# Using generalised dissimilarity modelling and targeted field surveys to gap-fill an ecosystem surveillance network

**DOI:** 10.1101/2020.06.01.107391

**Authors:** Greg R. Guerin, Kristen J. Williams, Emrys Leitch, Andrew J. Lowe, Ben Sparrow

**Affiliations:** School of Biological Science, The University of Adelaide, Adelaide, South Australia 5005, Australia; Terrestrial Ecosystem Research Network; CSIRO Land and Water, Canberra, Australian Capital Territory 2601, Australia

## Abstract

1. When considering which sites or land parcels complement existing conservation or monitoring networks, there are many strategies for optimising ecological coverage in the absence of ground observations. However, such optimisation is often implemented theoretically in conservation prioritisation frameworks and real-world implementation is rarely assessed, particularly for networks of monitoring sites.
2. We assessed the performance of adding new survey sites informed by predictive modelling in gap-filling the ecological coverage of the Terrestrial Ecosystem Research Network’s (TERN) continental network of ecosystem surveillance plots, Ausplots. Using plant cover observations from 531 sites, we constructed a generalised dissimilarity model (GDM) in which species composition was predicted by environmental parameters. We combined predicted nearest-neighbour ecological distances for locations across Australia with practical considerations to select regions for gap-filling surveys of 181 new plots across 18 trips. We tracked the drop in mean nearest-neighbour distances in GDM space, and increases in the actual sampling of ecological space through cumulative multivariate dispersion.
3. GDM explained 34% of deviance in species compositional turnover and retained geographic distance, soil P, aridity, actual evapotranspiration and rainfall seasonality among 17 significant predictors.
4. Key bioregions highlighted as gaps included Cape York Peninsula, Brigalow Belt South, South Eastern Queensland, Gascoyne and Dampierland.
5. We targeted identified gap regions for surveys in addition to opportunistic or project-based gap-filling over two years. Approximately 20% of the land area of Australia received increased servicing of biological representation, corresponding to a drop in mean nearest-neighbour ecological distances from 0.38 to 0.33 in units of compositional dissimilarity. The gain in sampled ecological space was 172% that from the previous 181 plots. Notable gaps were filled in northern and south-east Queensland, north-east New South Wales and northern Western Australia.
6. Biological scaling of environmental variables through GDM supports practical sampling decisions for ecosystem monitoring networks. Optimising putative survey locations via ecological distance to a nearest neighbour rather than to all existing sites is useful when the aim is to increase representation of habitats rather than sampling evenness *per se*. Iterations between modelled gaps and field campaigns provide a pragmatic compromise between theoretical optima and real-world decision-making.

## Introduction

Conservation planners have long grappled with the problem of how to maximise ecological representation in the absence of perfect information on biodiversity (Albuquerque & Beier 2018). For example, finite budgets mean the acquisition of incompletely surveyed land parcels to complement reserve systems must be prioritised. Indeed, much of the literature concerning spatial representation deals with conservation planning (Ferrier 2002; Williams et al. 2006; Arponen et al. 2008; Pennifold et al. 2017), and, in particular, reserve selection (Faith & Walker 1996; Araújo et al. 2001; Engelbrecht et al. 2016).

Fewer studies have addressed optimising representation for targeted biological surveys (Funk et al. 2005; Hortal & Lobo 2005; Ware et al. 2018) or the design of monitoring networks (Stohlgren et al. 2011; Rose et al. 2016). Networks of ecosystem observation plots have traditionally been designed to maximise spatial or environmental coverage using stratification (Goring et al. 2016; Guerin et al. 2020). Even so, the problems of complementary reserve selection and gap-filling monitoring networks are, in essence, the same (Hopkins & Nunn 2010); both seek to maximise ecological coverage in the absence of fine-scale biodiversity survey data. For this reason, biodiversity surrogates are often employed to estimate and compare representativeness (Grantham et al. 2010).

While many easily measured indicators of biological pattern could serve as surrogates, spatial information is most commonly employed, due to its relative ease of access and high spatial coverage, and to enable interpolation between sparse data locations (Rodrigues & Brook 2007). For example, environmental layers relating to soil type, climate and land cover are frequently used as surrogates for ecological community composition, as are maps of classified vegetation types or bioregions (Ware et al. 2018). The rationale for environmental layers as surrogates is the assumption that species compositional turnover occurs as a result of environmental heterogeneity, due to differences in realised species niches (Araújo et al. 2001). However, while environmental layers represent the broad environmental setting of ecological communities, they do not directly capture other key drivers of finer scale species compositional turnover, such as local disturbance regimes, microclimates, human impacts, and biotic interactions (Araújo et al. 2001; Peres-Neto et al. 2012; Goring et al. 2016). The spatial structure of ecological similarity (i.e., the degree to which species are shared between locations in space or time) is well known and is the reason that unstratified, systematic sampling performs reasonably at sampling ecological variation (Peres-Neto et al. 2012; Goring et al. 2016; Guerin et al. 2020). While local (patch level) drivers are opaque at large scales, ideally the purely spatial component of turnover would be included in assessments of surrogates.

In terms of selecting additional sites that best complement the ecological pattern of an existing set of sites, a range of optimisation strategies have been proposed that vary in terms of effectiveness and computational efficiency. Stohlgren et al. (2011) used Maxent modelling of the environmental coverage of monitoring site locations to select the most dissimilar putative sampling sites. Another approach is to select new sites that maximise the pairwise distances among all sample sites, so-called ‘Environmental Diversity’, which results in even sampling across environmental space (Faith & Walker 1996; Faith et al. 2004; Albuquerque & Beier 2018).

One realisation of environmental surrogate approaches is that ecological turnover is rarely a linear function of environmental turnover. The relationship is often nonstationary, with varying rates of turnover or even abrupt boundaries between compositionally distinct communities or geographic ecotones between structural vegetation types (Ferrier et al. 2007; Gibson et al. 2015; Goring et al. 2016). Generalised dissimilarity modelling (GDM; Ferrier et al 2007) has become the standard in community ecology for deriving biotically-scaled environmental surrogates (ecological environments) where sufficient biological training data are available (Ashcroft et al. 2010; Rose et al. 2016), and is an appropriate surrogate measure for Faith’s ED (Faith et al. 2004; Faith 2011). Because GDM explicitly accounts for nonlinearity, it more realistically scales environmental heterogeneity (and geographic distance) to actual species compositional turnover (Ferrier 2002; Ware et al. 2018). In doing so, GDM conveniently predicts ecological dissimilarity (and therefore the inverse measure, ecological similarity). The biotically-scaled environmental variables then characterise the pattern of ecological environments (Williams et al. 2014).

Here, we combined the predictive power of GDM with practical considerations to strategically gap-fill an established network of ecosystem surveillance plots – TERN Ausplots (Sparrow et al. 2019a). Ausplots were originally stratified to sample environmentally distinct bioregions across the Australian rangelands (Sparrow et al. 2019b; Guerin et al. 2020). However, due to an expansion in scope to all terrestrial bioregions and a practical limitation on the total number of sites that can be feasibly monitored, we aimed to select regions for survey that would efficiently increase ecological representation.

Because we aimed for representation and inclusion rather than even sampling across ecosystems (Schröder et al. 2006), we used a ‘nearest-neighbour’ approach (Faith & Norris 1989), in which each ecosystem is ideally ‘serviced’ by a representative site, with a preference to add new sites in habitats that are currently the least well represented (Hargrove & Hoffman 2004). We did not aim to make sampling even across all environments, not least because part of the original rationale behind Ausplots was that the Australian rangelands were more poorly monitored than mesic coastal areas (Guerin et al. 2017).

Here, we present how GDM was used to guide the placement of gap-filling surveys and assess the performance of strategic field sampling over time to fill identified gaps and increase the diversity (but not necessarily evenness) of the sampling in environmental space. While most of the literature on this topic selects putative, optimal sites (Arponen et al. 2008; Hoffman et al. 2013; Rose et al. 2016; Albuquerque & Beier 2018), it is because we implemented the approach via field surveys that we use real new sites and observations of species turnover to assess performance. This assessment of biological survey gaps in the TERN Auplots network was guided by the following questions:

### Generalised dissimilarity model

– What are the main macro-scale predictors of plant species turnover across Australia?
– Is extrapolation needed to predict turnover at national scale?

### Site selection

– Surveys of which putative regions would ecologically complement the existing Ausplots network, based on GDM predictions?
– How well did gap-filling surveys ‘service’ under-sampled ecosystems in practice?

## Methods

### Ausplots method and data

TERN Ausplots are a distributed network of one-hectare, fixed location monitoring plots with scope covering all major terrestrial ecosystems in Australia. Field methods (described at length elsewhere) focus on sampling vegetation and soils (White et al. 2012; Sparrow et al. 2019b). We calculated percent cover of plant species from 1010 point-intercepts per plot, with identifications of all species observed anywhere in the plot made through herbarium determination of vouchers. We accessed data for 763 Ausplots, including 582 plots that had been established prior to commencement of our gap-filling objectives (Guerin et al. 2018; TERN 2020). For convenience, we refer to the first 582 plots hereafter as ‘establishment plots’ and the subsequent 181 plots surveyed over 18 trips as ‘gap plots’.

### Generalised dissimilarity modelling

We fit a GDM where the response variable was compostional turnover in plant species among establishment plots using species percentage cover as the abundance measure in calculating Bray-Curtis dissimilarity (Taft 2014; Manion et al. 2017; R Core Team 2016). We included repeat surveys (representing 7% of the samples) to maximise ecological information content, resulting in 572 surveys from 531 unique plots that had full vegetation data available at that time. Revisits capture observed variation in species composition within the same environmental setting and are handled numerically through spatial covariables, in that temporal replicates will have zero spatial distance.

Spatial layers with a resolution of 9 arcseconds were used as predictors, comprising 44 candidate variables related to climate, soil and landscape (Grundy et al. 2015; Harwood et al. 2016; Gallant et al. 2018; Table 1). Geographic distance was also included as a predictor. The number of predictors was reduced to a subset of 25 for which variance inflation factors were less than 10 by removing highly correlated variables that had weak importance in exploratory models. A statistical variable selection process for model fitting was applied to these 25 candidate variables.

**Table 1.** Climate, soil and landform variables used in a generalised dissimilarity model (GDM) of plant species composition based on TERN Ausplots ‘establishment’ sites (source: Harwood et al. 2016; Gallant et al. 2018). Variables ranked by importance according to summed model coefficients (height of the *I*-spline on the y-axis as a measure of relative importance).

| Code | Name and origin data citation | Unit | Summed GDM coefficients |
| --- | --- | --- | --- |
| GEO | Geographic distance | degrees | 2.72 |
| PTO | Total Phosphorus (Viscarra Rossel et al. 2014b) | % | 2.22 |
| EAA | Annual total actual evapotranspiration (Harwood et al. 2016) | mm | 2.10 |
| ADI | Minimum monthly aridity index (Williams et al. 2012) | proportion | 1.74 |
| ADX | Maximum monthly aridity index (Williams et al. 2012) | proportion | 1.45 |
| PTS2 | Precipitation seasonality 2 (Williams et al. 2012) | ratio | 1.36 |
| TRX | Maximum monthly mean diurnal temperature range (Harwood et al. 2016) | °C | 1.02 |
| NTO | Total Nitrogen (Viscarra Rossel et al. 2014c) | % | 0.88 |
| PHC | pH – CaCl2 (Viscarra Rossel et al. 2014d) | None | 0.78 |
| BDW | Bulk Density – Whole Earth (Viscarra Rossel et al. 2014e) | g/cm3 | 0.73 |
| CLY | Clay (Viscarra Rossel et al. 2014f) | % | 0.60 |
| AWC | Available Water Capacity (Viscarra Rossel et al. 2014g) | % | 0.57 |
| PTS1 | Precipitation seasonality 1 (Williams et al. 2012) | ratio | 0.47 |
| SOC | Organic Carbon (Viscarra Rossel et al. 2014h) | % | 0.38 |
| EPX | Maximum monthly potential evaporation (Harwood et al. 2016) | mm | 0.26 |
| CONAREA | Contributing area (Gallant & Austin 2012) | index | 0.11 |
| ECE | Effective Cation Exchange Capacity (Viscarra Rossel et al. 2014i) | meq/100g | 0.04 |

To build the model, we used a backward selection procedure. Variables were removed when model coefficients summed to zero, which indicates no effect on species turnover. Variables were also removed if they were not statistically significant, using the method of Leitão et al. (2017), applying 1000 matrix permutations to test whether the contribution of each variable in the GDM was greater than expected at random.

### Selection and appraisal of gap-filling surveys

Gaps in the ecological coverage of establishment plots were assessed spatially. Using predictor layers that had been GDM-transformed according to corresponding modelled *I*-splines (Manion 2009), we calculated the Euclidean distance of each grid cell to the nearest establishment plot in GDM space, and produced heat maps of ‘ecological distance to nearest neighbour’ (NND), which spatially maps representativeness (Hoffman et al. 2013). To evaluate reliability, we mapped a model extrapolation index, calculated as the sum of GDM-transformed values outside the input data range (i.e., the total amount of extrapolated turnover; Gibson et al. 2015).

Various procedures can be used to optimise the selection of putative sites with the aim of reducing mean NND, indicating better ‘servicing’ by the plot network (Faith & Norris 1989). For example, in the *p*-median approach, the mean drop in NND across a set of demand points is optimised (Engelbrecht et al. 2016). A simpler ‘greedy’ approach is to sequentially select sites with the highest NND, though this does not optimise the servicing of other sites *per se*. In the real world, however, the selection of sites for surveys combines quantitative information with a set of practical considerations. Field expeditions are planned around appropriate seasonal conditions for floristic sampling, logistical constraints (such as time in the field, availability of road infrastructure) and access to suitable sampling locations. Using modelled information on gaps from the GDM, we targeted regions rather than specific sites. During these field trips, an average of 10 (range 2–16) plots were established to cover local variation and provide some level of replication. In some instances, gap surveys were combined with local spatial gap-filling around establishment sites, which were opportunistically conducted during surveys focused on revisits, or were established in collaboration with other ecosystem monitoring or surveillance survey programs.

To improve confidence in decision-making, we supplemented the modelling and NND heat maps presented here to gaps highlighted using a pre-existing GDM of plant species composition based on natural history records (‘ANHAT’ data; Williams et al. 2013; Ware et al. 2018). By applying the same set of establishment sites, *p*-median was used to map the highest priority areas for gap surveys (using the ‘Survey Gap Analysis Tool’; Funk et al. 2005; Manion & Ridges 2009). Confidence in the mapping of gaps was raised where the two sources of modelled gap information (i.e., GDMs based on Ausplots or ANHAT data) were similar. We do not report this existing modelling in full here as it has been described elsewhere and formed only one aspect of the decision-making process. Notwithstanding, the major gaps identified by either model were similar.

The identification and surveying of gaps continued iteratively. Here, we evaluate two years of gap surveys in two ways. Firstly, we add real field plots clustered in regional ‘trips’ to the establishment plots in the chronological order they were surveyed and calculate the drop in NND across Australia. We refitted the GDM and repeated the appraisal of gaps incorporating the additional gap-filling survey data. The results were highly similar in terms of responses to gradients, and spatial mapping of predicted remaining gaps (qualitatively identical), hence are not reported here in further detail.

Secondly, we use the species composition percentage cover data recorded during the field surveys to calculate cumulative multivariate dispersion (MVD) in ecological space (Bray-Curtis dissimilarities in species composition), as individual plots were surveyed in chronological order. MVD is defined as mean distance of plots to their centroid in Principal Coordinates space (MDS of pairwise site distances; Oksanen et al. 2018), and is a measure of beta diversity (Anderson et al. 2006). If gap-filling surveys have been successful, we would expect NND to decrease over time, while we would expect MVD to increase. The difference between successive NND heat maps visualises gap-filling spatially.

## Results

The GDM included geographic distance and 16 environmental variables as predictors, and explained 34% of deviance in pairwise dissimilarity (Table 1; Figs 1–2). The highest turnover was predicted in response to geographic distance, followed by soil P, aridity, actual evapotranspiration and rainfall seasonality. The GDM extrapolation index was below the suggested maximum of 0.1 (Gibson et al. 2015) for all but a small area in South Australia that was not highlighted as a gap (Fig. 1b).

**Fig. 1.**
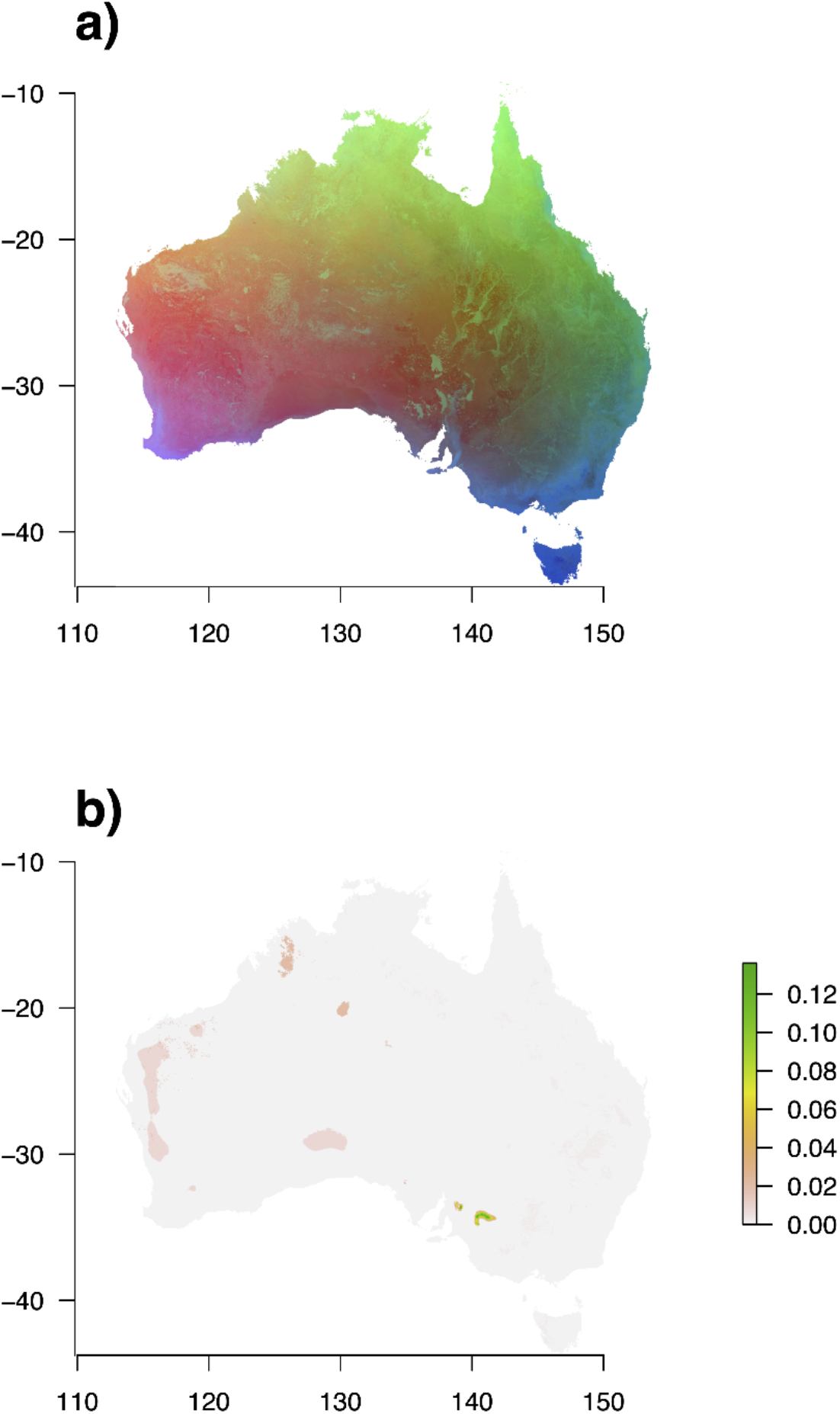
a) Visual representation of the generalised dissimilarity model (GDM) based on vascular plant species composition in TERN Ausplots ‘establishment’ plots. RGB colour scale represents position along the first three PCA axes of GDM-transformed predictors included in the fitted model. Similarity in colour represents similarity in species composition; b) GDM extrapolation index: sum of transformed grid values outside the input data range (total amount of extrapolated turnover).

**Fig. 2.**
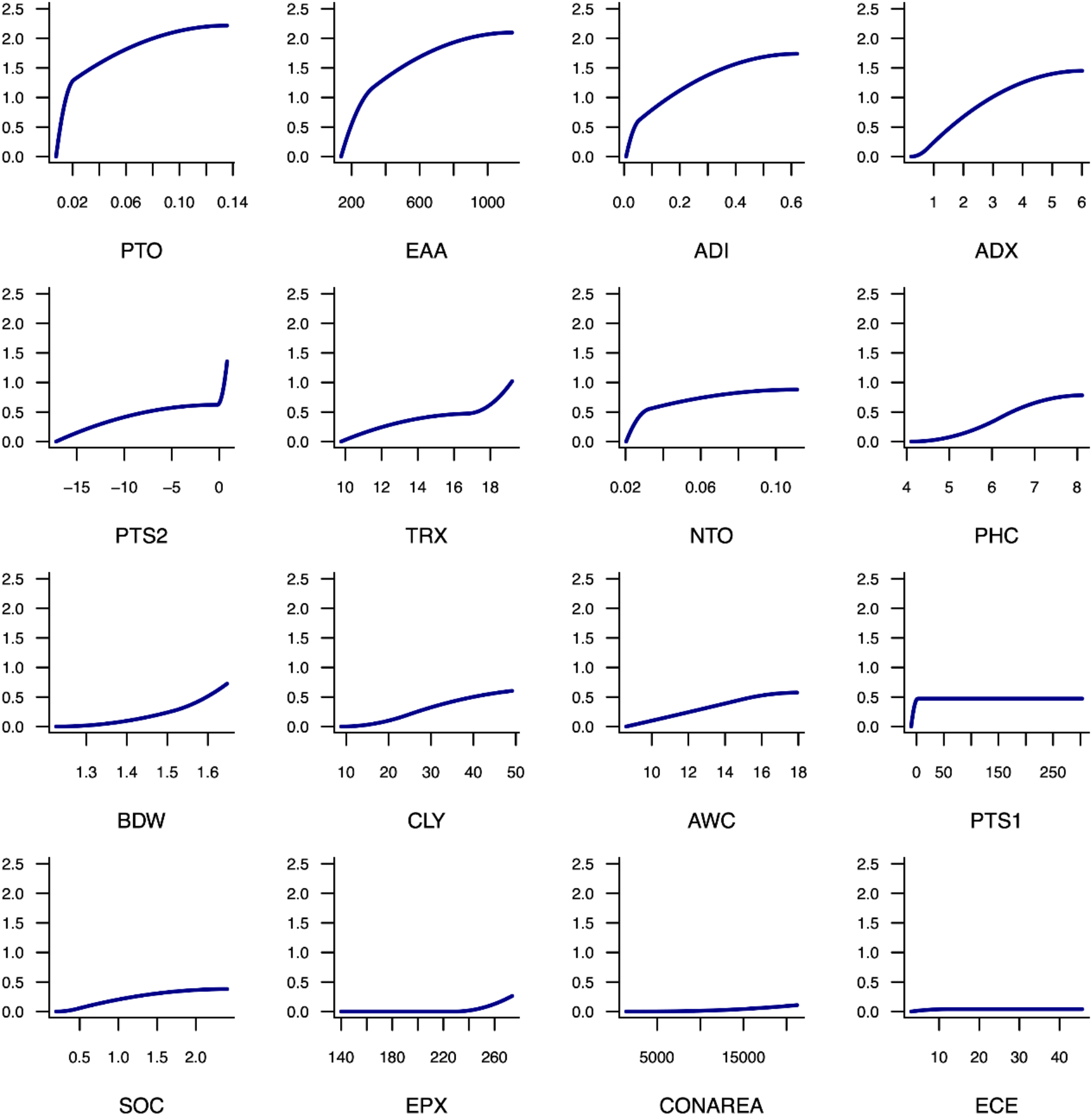
GDM *I*-spline partial functions, depicting the rate of species turnover along the 16 environmental gradients retained in the model, in order of variable importance, based on height of the *I*-spline function. See Table 1 for variable descriptions and importance values.

Prior to gap-filling surveys, the NND map highlighted the Cape York Peninsula (CYP), Brigalow Belt South (BBS) and South Eastern Queensland (SEQ) bioregions as notable gaps (Thackway & Creswell 1995; Figs 3a, 4). Additional gaps included the Gascoyne (GAS) and Dampierlands (DAL) bioregions, as well as the north-west deserts. Field campaigns over two years in 2018, 2019 and 2020 targeted these and other successive gaps regions. Eighteen trips, surveying a total of 181 plots, were conducted (Table 2; Figs 3b, 4).

**Table 2.**
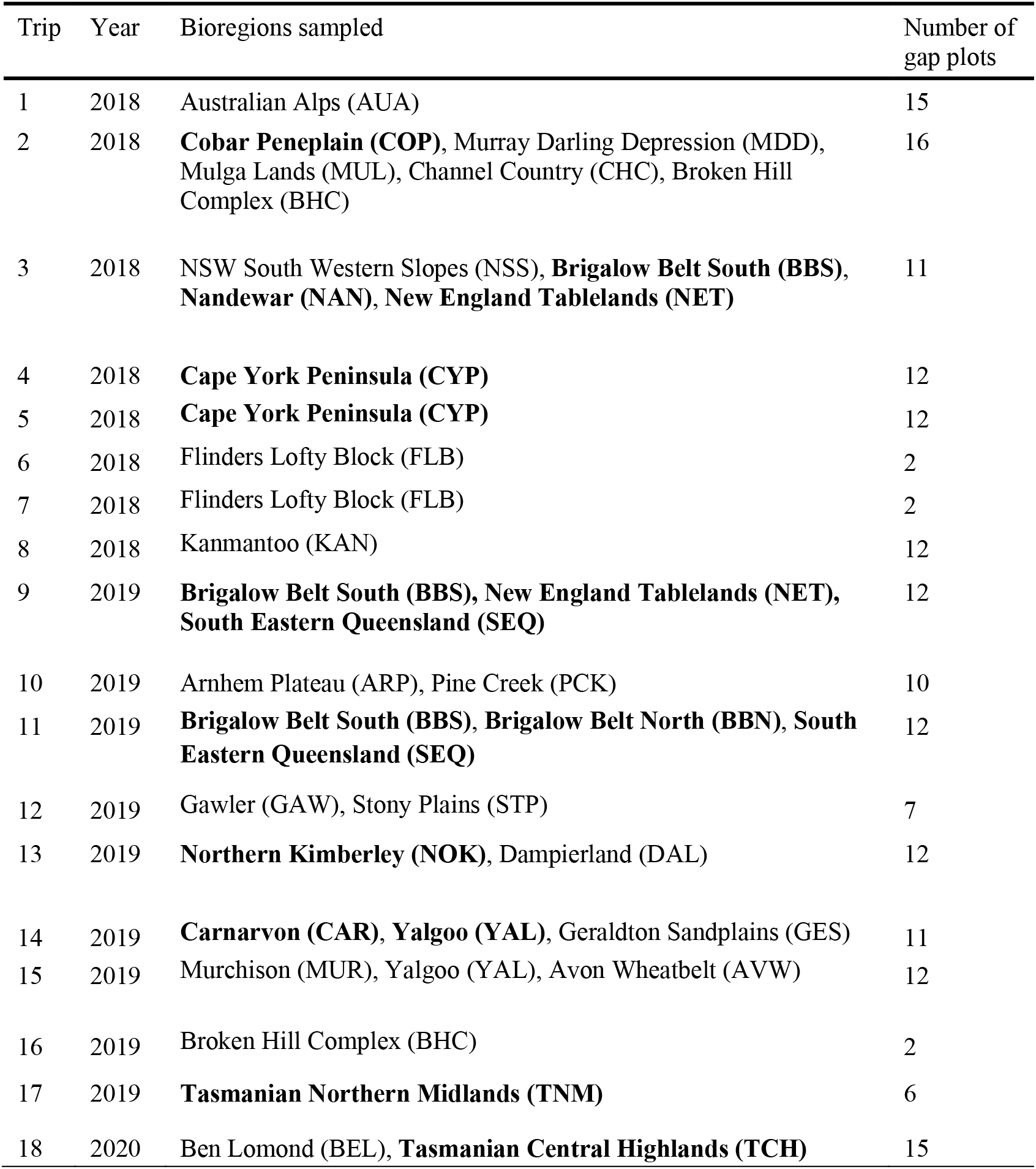
Survey trips during the gap-filling period of 2018–2020 in chronological order, noting the location and number of 181 new plots. Some trips also included revisits. Bold entries indicate bioregions (Fig. 4) that were not sampled prior to gaps surveys.

| Trip | Year | Bioregions sampled | Number of gap plots |
| --- | --- | --- | --- |
| 1 | 2018 | Australian Alps (AUA) | 15 |
| 2 | 2018 | **Cobar Peneplain (COP)**, Murray Darling Depression (MDD), Mulga Lands (MUL), Channel Country (CHC), Broken Hill Complex (BHC) | 16 |
| 3 | 2018 | NSW South Western Slopes (NSS), **Brigalow Belt South (BBS)**, **Nandewar (NAN)**, **New England Tablelands (NET)** | 11 |
| 4 | 2018 | **Cape York Peninsula (CYP)** | 12 |
| 5 | 2018 | **Cape York Peninsula (CYP)** | 12 |
| 6 | 2018 | Flinders Lofty Block (FLB) | 2 |
| 7 | 2018 | Flinders Lofty Block (FLB) | 2 |
| 8 | 2018 | Kanmantoo (KAN) | 12 |
| 9 | 2019 | **Brigalow Belt South (BBS)**, **New England Tablelands (NET)**, **South Eastern Queensland (SEQ)** | 12 |
| 10 | 2019 | Arnhem Plateau (ARP), Pine Creek (PCK) | 10 |
| 11 | 2019 | **Brigalow Belt South (BBS)**, **Brigalow Belt North (BBN)**, **South Eastern Queensland (SEQ)** | 12 |
| 12 | 2019 | Gawler (GAW), Stony Plains (STP) | 7 |
| 13 | 2019 | **Northern Kimberley (NOK)**, Dampierland (DAL) | 12 |
| 14 | 2019 | **Carnarvon (CAR)**, **Yalgoo (YAL)**, Geraldton Sandplains (GES) | 11 |
| 15 | 2019 | Murchison (MUR), Yalgoo (YAL), Avon Wheatbelt (AVW) | 12 |
| 16 | 2019 | Broken Hill Complex (BHC) | 2 |
| 17 | 2019 | **Tasmanian Northern Midlands (TNM)** | 6 |
| 18 | 2020 | Ben Lomond (BEL), **Tasmanian Central Highlands (TCH)** | 15 |

**Fig. 3.**
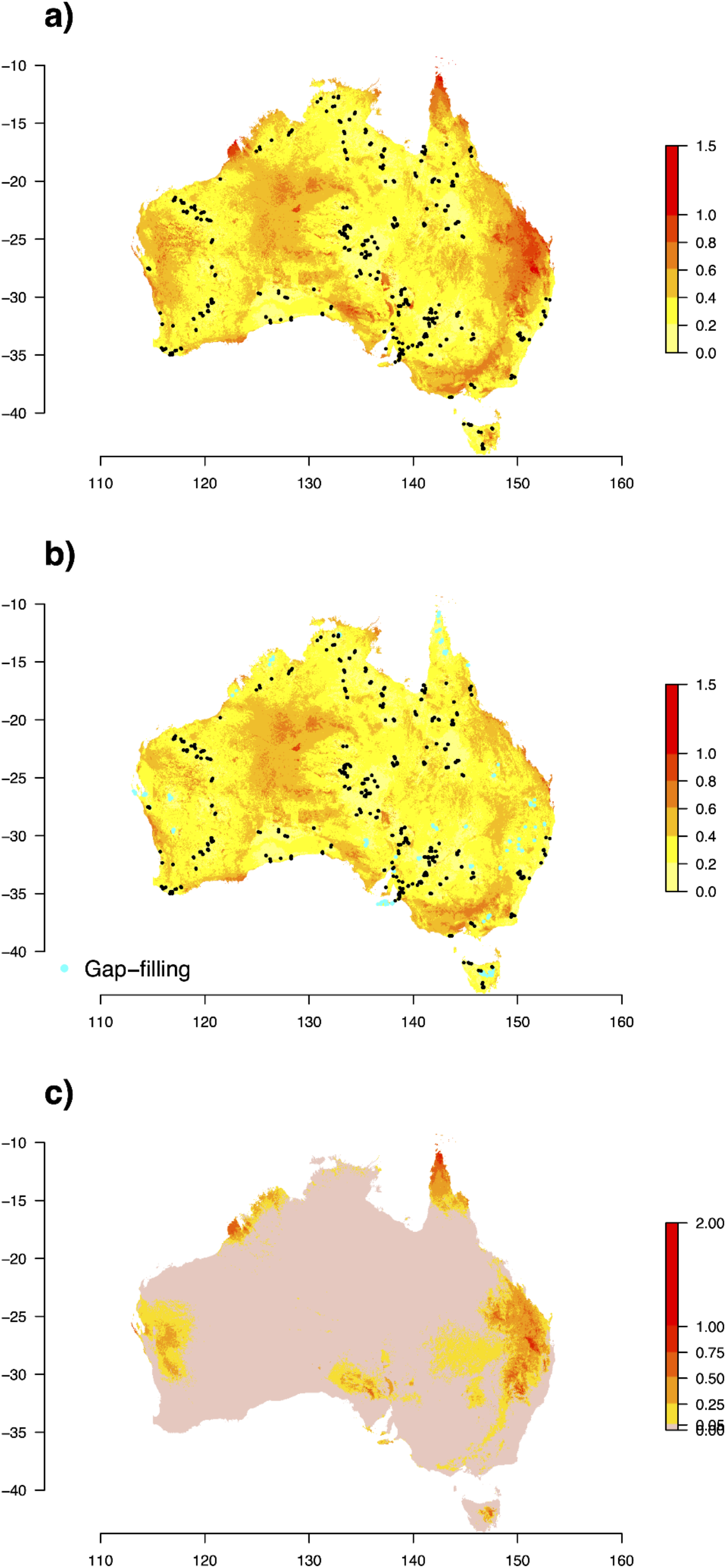
a) Map of Australia showing ecological distance (Euclidean distance in biotically-scaled environmental space) of each pixel to the most similar Ausplots site (NND), as predicted from a GDM of plant species composition. Based on 582 ‘establishment’ plots (points). High scoring regions represent ecological gaps in the sampling coverage that were targeted in subsequent surveys; b) NND as in previous panel, based on 763 plots (points), including 181 gap-filling plots; c) drop in ecological distance to most similar Ausplots location (NND) after 181 gap-filling plots were surveyed over 18 trips; calculated as the difference between maps before (a) and after (b) gap-filling.

**Fig. 4.**
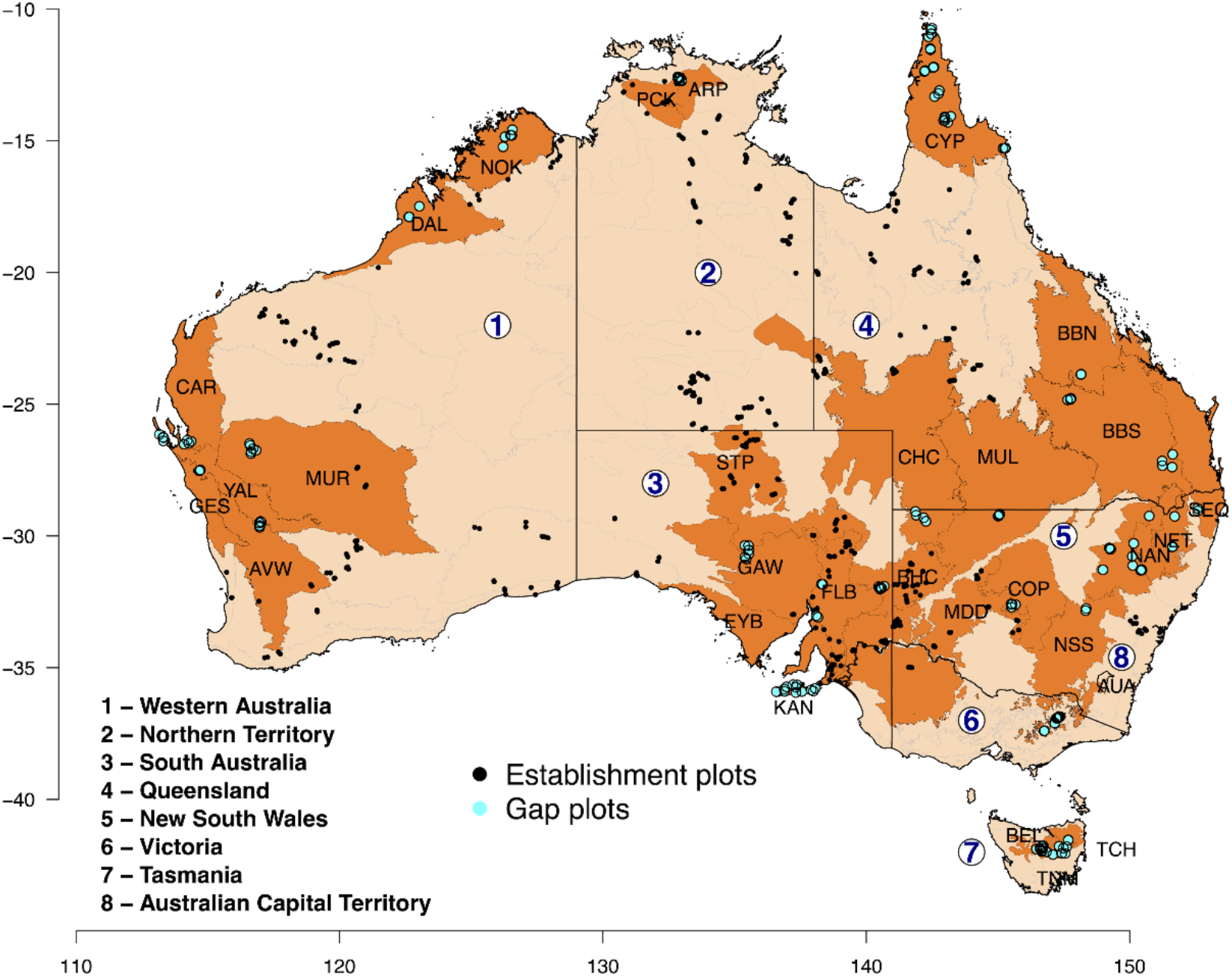
Australia, highlighting Australian States and Territories and the IBRA bioregions targeted by ‘gap plots’ in 2018–2020 (Table 2; Department of the Environment 2012).

Considering all surveyed locations as of 2020 and NND, the major remaining gap was the north-west desert region, and to a lesser extent the far south-east, the coast sandplains of south-west Western Australia and the southern Queensland coastline (Figs 3b; 4). The difference between before and after NND maps spatially highlights the gaps that were filled by the surveys (Fig. 3c). Many of the major gaps identified initially via NND were filled to some extent. The magnitude of the improvement in NND for those gap areas was high, with NND drops of up to ~1, equivalent to a shift from no coverage to complete servicing of species composition based on the GDM.

Mean NND across Australia dropped from 0.38 to 0.33, in units of compositional dissimilarity (Fig. 5a). The proportion of Australia that was serviced through gap-filling plots via decreased NND was 18% at a minimum of 0.05 drop, 14% at a minimum of 0.1 drop, 6% at a minimum drop of 0.25, and 1% at a minimum 0.5 drop (Fig. 5b).

**Fig. 5.**
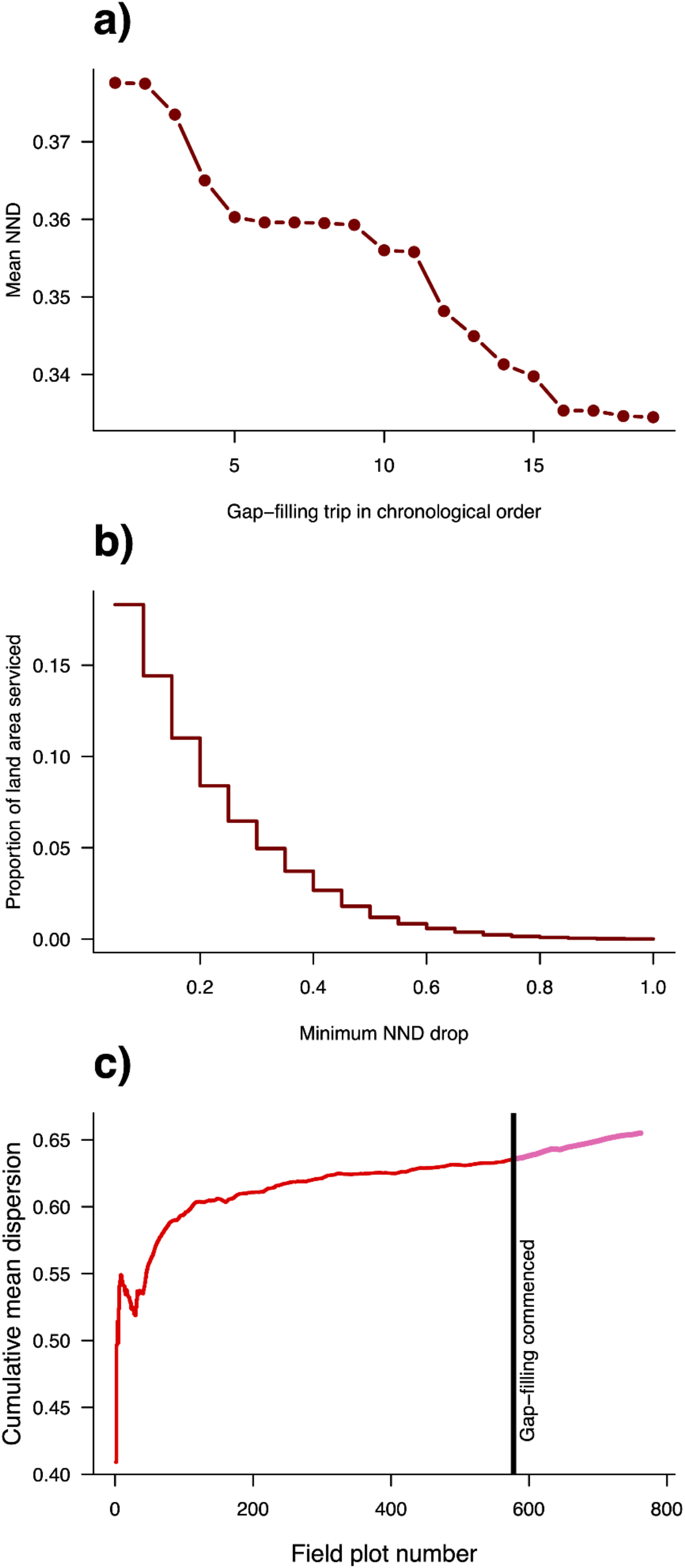
a) Drop in mean ecological distance to nearest sampled Ausplots site (NND) with the addition of 18 gap-filling trips in chronological order. First point is establishment plots mean. Trips were expeditions targeting particular regions in which an average of 10 (range 2–16) new plots were surveyed. Some trips focused on revisits to established sites, with smaller numbers of local gap-filling plots being established (e.g. 7^th^ and 8^th^ points above, corresponding to trips 6–7 in Table 2); b) Cumulative proportion of Australia’s land area serviced by decreased distance to nearest Ausplots site in ecological space (NND) due to 18 gap-filling surveys 2018–2020 across a range of minima. 18% of the continent was serviced with a minimum NND drop of 0.05; c) Changes in the real sampling of ecological space as visualised via cumulative mean multivariate dispersion (y-axis; MVD, mean distance to centroid in Principal Coordinates Space, based on MDS of Bray-Curtis dissimilarities). The x-axis represents additional surveys of plots in chronological order.

The rate at which mean MVD increased with additional surveys (representing accumulation of ecological space in the network) rose sharply during the early establishment phase of surveys, levelled off from approximately 200 plots, but was then higher during the gap-filling phase, with a gain of 172% that of the previous 181 plots, and an 8% increase in overall accumulation over the entire establishment phase (Fig. 5c).

## Discussion

Biodiversity surrogates provide a powerful approach to gap-filling when applied to networks of both conservation reserves and surveillance monitoring sites (Rodrigues & Brook 2007). The utility of environmental layers as surrogates increases when these are biotically scaled by modelling their correlation with species turnover (Ware et al. 2018). We chose generalised dissimilarity modelling (GDM) for that purpose because it accounts appropriately for nonlinear responses, including varying rates of turnover at different positions along environmental gradients (Ferrier 2002), and has been extensively applied and proven as suitable for biodiversity assessment with hundreds of citations (Ferrier et al. 2007).

We were able to use the comprehensive species composition and relative abundance data recorded in the large set of plots already established across Australia to inform further gap-filling surveys through compositional models. This approach contrasts with the original environmental stratification used to establish the TERN Ausplots network, which used environmental layers in the absence of ground data at that stage (Guerin et al. 2020).

Although species occurrence data held in natural history databases provide greater spatial and taxonomic coverage, the advantage of TERN Ausplots species composition data for modelling ecological distance is its ability to capture locally complete sets of co-occurring species at patch scales, and is the only standardised dataset of species identity and relative abundance within one-hectare plots across all State and Territory jurisdictions in Australia. Alternative datasets for this modelling are either presence-only (requiring assumptions about aggregation to represent assemblages) or require harmonisation of plot-based surveys with eclectic methods (e.g., plot sizes, affecting species richness) and different accuracies for scoring relative abundance, necessitating development of covariates to characterise and explain systematic error.

By mapping predicted ecological distance to the most similar site in the establishment network of permanent plots, we were able to identify major sampling gaps. These gaps were addressed via survey expeditions to the Australian Alps (AUA), Cape York Peninsula (CYP), Brigalow Belt South (BBS), Yalgoo (YAL), Dampierland (DAL), Tasmanian Northern Midlands (TNM) and Tasmanian Central Highlands (TCH) bioregions (Table 2; Fig. 4; Thackway & Creswell 1995). Our retrospective assessment confirms that the combination of gap modelling and practical considerations with more traditional local stratification of survey plots was able to achieve the goal of efficiently representing terrestrial Australian ecosystems.

### Predictors of plant species turnover across Australia

In the GDM of species composition data from Ausplots, the highest species turnover was fitted to geographic distance, soil P, aridity (monthly minimum and maximum aridity index, potential evaporation and actual evapotranspiration) and rainfall seasonality. However, our aim was to model ecological turnover rather than determine the most important drivers, and it is likely that covariance among competing predictors is involved in some of the observed patterns. Indeed, variance partitioning (Gilbert & Bennet 2010) shows total deviance explained comprised 3.8% independent spatial effect, 34.6% independent environmental effect and 61.7% spatial–environmental covariance, while the partitioning for environmental variables was 31.9% independent climate effect, 21.3% independent soil effect and 46.9% climate–soil covariance.

The deviance explained by the model at 34% suggests that a number of additional factors not accounted for in the model influence observed species composition, as expected. Examples of such factors include fire and grazing regimes, human disturbance, proximity to regional climate refugia, seasonal phenology and climatic condition, and biotic interactions (Araújo et al. 2001; Gibson et al. 2015; Keppel et al. 2017), and these are likely to weaken relationships between macroclimate and biodiversity, particularly at local scales (Bruelheide et al. 2018; Harrison et al. 2020).

The other published national-scale model of turnover in plant species composition for Australia is the GDM based on occurrence data extracted from the Australian Natural Heritage Assessment Tool (‘ANHAT’; Williams et al. 2013; Ware et al. 2018). The ANHAT model was also fitted with a backwards selection procedure that finished with 11 climate and six substrate predictors. The ANHAT and Ausplots GDMs are similar, with rainfall seasonality, evaporation and actual evapotranspiration as the most important predictors. The importance of aridity and rainfall seasonality are intuitive as key variables at large scales that explain differences between tropical, desert and temperate biomes (as visualised in Fig. 1a). However, it is likely that the importance of particular variables, and the nature of the ecological response to them, varies from region to region (Burley et al. 2012; Guerin et al. 2019).

Edaphic predictors are harder to compare between the two models because the ANHAT model used an earlier version of the soil parameters before the full TERN data release (Viscarra-Rossel & Chen 2011; Viscarra Rossel et al. 2014a; Grundy et al. 2015; Viscarra Rossel et al. 2015), whereas our Ausplots model used a more comprehensive variable suite aggregated to 9-arcsecond grids from the 3-arcsecond source data (Viscarra Rossel et al. 2014a; Gallant et al. 2018). The ANHAT GDM incorporates a larger dataset of presence-only records aggregated within adequately sampled grid cells (as described in Williams et al. 2010), whereas the Ausplots GDM was based on comprehensive floristic surveys of 531 plots, with full inventories of co-occurring species at patch level along with robust measures of percent cover.

### Complementing the existing TERN Ausplots network with strategic gap-filling surveys

Strategic gap-filling surveys guided by the Ausplots GDM and practical considerations were successful in complementing sampling of the Australian vegetation over two years. The mean nearest-neighbour distance (NND) across the whole of Australia dropped from 0.38 to 0.33 in units of compositional dissimilarity. Almost one fifth of the continent was thereby additionally serviced by gap-filling to some degree, while 6% of the land area of Australia had a drop in NND of at least 0.25 in units of compositional dissimilarity. The size of the additional ecological space sampled was 172% that of the previous 181 plots, adding 8% over all establishment phase sampling, as measured by multivariate dispersion, which is significant given the law of diminishing returns and that coverage of the Australian terrestrial environment by establishment plots was already quite comprehensive (Fig. 5c; Guerin et al. 2020).

Although we generated sets of putative survey sites optimised in one way or another during the course of the gap-filling process, we found it more enlightening to evaluate how gap-filling performed in practice. While optimisation methods are useful, it is not always feasible to access pre-defined locations. Logistical considerations also influence survey locations. The *modus operandi* of the TERN ecosystem surveillance field team is to target a region with a set of surveys over an expedition for cost-efficiency reasons, to capture local heterogeneity and to ensure a level of replication.

Planning of field surveys is iterative and often involves a series of logistical contingencies. To achieve all of this, an optimal approach would need to iteratively trade-off survey ‘benefit’ due to greater biological coverage against survey ‘cost’ (e.g. Rodewald et al. 2019). In covering the diverse ecosystems of Australia, field work needed to be flexible, practical and pragmatic to maximise useful data collection given huge variation in seasonal climatic conditions and plant phenology responses. In addition, opportunities for sampling due to collaborative projects and the time and cost of reaching distant and remote sampling locations, were among many factors that must be considered. Sampling of the diverse Australian environment has been remarkably efficient for a relatively small number of plots, due to strategic selection of monitoring sites.

### The never-ending story

Strategic field surveying of identified gaps is an iterative process leading to gap-filling at finer levels of detail over time. A practical end-point to this process is imposed by resource constraints that limit the number of field plots feasible to establish and regularly revisit, considered against the desired level of gap-filling and trade-offs between spatial and temporal sampling coverage. The goal of TERN Ausplots has been to establish a representative network of ecosystem surveillance plots to remeasure at least once per decade. Gap-filling will continue to some degree into the future, with increasing emphasis on revisits.

In terms of the GDM developed here, a feasible end-point could be to continue gap-filling until all NND are below one (see Fig. 3), meaning each grid cell is predicted to share some species with the plot network, based on the most up-to-date data available (Ferrier 2002). This end-point has almost been reached already, with only five 100,000ths of the land area over that cut-off. A more ambitious end-point would be a maximum NND of 0.5. This end-point requires increased servicing of an additional 12% of the Australian land area (a reduction from 21% prior to gap-filling; Fig. 3). Current indications are that a network with an additional 250–750 plots will be adequate to cover the major Australian terrestrial environments with sufficient replication of major vegetation types.

The largest remaining gap identified in the TERN Ausplots network from the GDM is a large region in the north-west deserts (Fig. 2b). This area was already an obvious spatial gap in the distribution of field plots. However, staging an expedition to this region with the expectation of performing revisits requires careful, long-term planning, as it is a remote and challenging environment far from population centres and with extremely limited road infrastructure. In addition, long-term access to sites for monitoring would need to be secured through partnerships with land managers and traditional owners which we aim to progress in the coming years.

The never-ending pursuit of ecological sampling gaps in the ecosystem surveillance network also continually shifts priority sites for regular revisits. For example, the subset of Ausplots that is most representative of the network, and of Australian ecosystems collectively, will evolve as gap-filling surveys proceed. This iterative approach to representation is one consideration in sequencing repeat visits. The existence of climate change and other drivers of compositional dynamics is another consideration. Gaps in representativeness may therefore change at different rates across the region. For this reason, sensitivity and exposure to climate change are now also being considered empirically to ensure ecosystems potentially undergoing rapid change are more regularly resampled.

### Limitations

While we implemented a quantitative yet practical solution to ecological gap-filling surveys, a range of other factors will also determine the efficacy of the network in observing spatial and temporal patterns in Australian terrestrial ecosystems. The frequency of revisits, precision of measurements relative to rates of change and degree of replication are factors that contribute to the power to detect compositional change in species and infer drivers of observed patterns (Guillera-Arroita & Lahoz-Monfort 2012). Remotely derived environmental layers, even when biotically scaled, are imperfect surrogates for biodiversity turnover (Araújo et al. 2001) for reasons already discussed. Other aspects of ecological stratification and gap-filling could include sampling along gradients of disturbance and human influence, for example.

### Conclusions

Gap-filling surveys for ecosystem monitoring or conservation networks are guided by quantitative assessments that help to prioritise putative target sites within practical constraints. Ecological coverage of TERN Ausplots was significantly increased by targeted surveys guided by biotically scaled environmental surrogates for biodiversity. Optimising putative survey locations by modelling distance to a nearest neighbour in GDM space was an efficient analytical method that enabled regular updates in near real time. Filling ever-finer gaps becomes a trade-off with temporal coverage as well as addressing different kinds of gaps, such as relative climate change vulnerability and consideration of logistical constraints. Modelling combined with field campaigns in near real time offers a pragmatic compromise for real-world decision-making.

## Acknowledgements

We thank all members past and present of the TERN Ecosystem Surveillance field and data teams and TERN, supported by the Australian Government through the National Collaborative Research Infrastructure Strategy.

## Notes

### Competing Interest Statement

The authors have declared no competing interest.

